# Barrier Function of the Vero E6 Cell Monolayer

**DOI:** 10.1101/2025.09.19.677358

**Authors:** Julia Tarnoki-Zach, Elod Mehes, Bernadett Palyi, Szilvia Bosze, Andras Czirok

## Abstract

Vero E6 cells are a standard model in virology research because of their broad susceptibility to viral infection. However, their ability to form functional barrier layers has received less attention. Here, we show that Vero E6 cells cultured on transwell inserts develop continuous monolayers with moderate transepithelial electrical resistance (TEER), reaching approximately 30 Ω·cm^2^ in serum-containing medium and 45 Ω·cm^2^ in virus production medium. The permeability coefficient (Papp) for 4 kDa FITC-dextran, a common measure for paracellular transport, is 3.7 × 10^−6^ cm/s, comparable to established endothelial models such as HUVEC or corneal endothelium. Immunofluorescence confirms the presence of tight junction (ZO-1) and adherens junction (β-catenin) proteins. Chelation of calcium with EGTA causes a dose-dependent, reversible decrease in TEER and a marked increase in permeability, confirming the calcium sensitivity of the barrier. These findings suggest that Vero E6 cells serve as a valuable epithelial barrier model, facilitating studies on viral entry, immune evasion, and drug delivery.

## 1 Introduction

The integrity and selectivity of biological barriers, such as epithelial and endothelial layers, are required for compartmentalization and homeostasis in multicellular organisms Yeste et al. [2018], Lechuga et al. [2024]. *In vitro* models of these barriers are used to study transport processes, immune signaling, and host–pathogen interactions Helms et al. [2016], Chaulagain et al. [2023]. Characterization typically relies on trans-epithelial electrical resistance (TEER), macromolecular permeability, and the organization of junctional proteins Srinivasan et al. [2015], Bednarek [2022].

Many epithelial cell lines are used as barrier models, with MDCK Misfeldt et al. [1976], Irvine et al. [1999] and Caco-2 Hidalgo et al. [1989] cells being the most notable. However, barrier characterization has not been the main focus for other cell lines widely used in research. The Vero E6 (also known as VERO C1008) line Mizusawa et al. [1988], Terasima et al. [1988], Earley and Johnson [1988], Ammerman et al. [2008], derived from African green monkey *C. sabaeus* kidney is extensively used in SARS-CoV-2 and other virology research Ogando et al. [2020], Essaidi-Laziosi et al. [2021], Souza et al. [2022], Tóth et al. [2023]. Vero E6 cells are preferred for their stable growth characteristics and susceptibility to infection, owing in part to deficient interferon production Emeny and Morgan [1979], Konishi et al. [2022], Prescott et al. [2010]. Their epithelial origin, cobblestone morphology, and expression of junctional proteins suggest a potential to serve as a functional barrier model, which is especially relevant when researching viruses that target epithelial surfaces.

Establishing Vero E6 as a barrier model provides a unique opportunity to link virology and barrier biology. Many viruses, including coronaviruses, influenza viruses, flaviviruses, and enteroviruses, infect hosts by crossing epithelial or endothelial barriers, disrupting tight junctions or paracellular permeability to reach deeper tissues Short et al. [2016], Tugizov [2021], Ding et al. [2022], Linfield et al. [2021]. A model system that combines viral permissiveness and measurable barrier properties would allow for direct studies of how viral replication interacts with epithelial integrity. Unlike traditional barrier models, Vero E6 cells are already widely used in virology laboratories and have well-documented infection kinetics, reducing the need for additional culture systems. This dual utility makes them an attractive platform for mechanistic studies and compound screening in contexts where barrier function and viral infection intersect.

Here, we investigate the barrier properties of Vero E6 monolayers under standard conditions. Specifically, we characterize the temporal development of TEER, assess macro-molecular permeability using 4 kDa FITC-dextran, and test the reversibility of barrier integrity after EGTA-mediated calcium chelation. Together, these experiments establish Vero E6 as a low-resistance but functional epithelial barrier model with particular value for infection-related studies.

## 2 Materials and Methods

### 2.1 Reagents, buffers and media

Dulbecco’s Modified Eagle’s Medium (DMEM) with or without phenol red, phosphate buffered saline (PBS), and L-glutamine were obtaied from Lonza (Basel, Switzerland). Sodium pyruvate and trypsin were purchased from Sigma-Aldrich (St Louis, MO, USA). Non-essential amino acids, fetal bovine serum (FBS) and penicillin/streptomycin (10,000 units penicillin and 10 mg streptomycin/mL) were obtained from Gibco (Thermo Fisher Scientific, Waltham, MA, USA).

To maintain cell cultures, time-lapse imaging and for growth on transwell inserts, DMEM with phenol red was supplemented with 10% FBS, 2 mM L-glutamine, 100 µg/mL penicillin/streptomycin, 1 mM sodium piruvate and 1% non-essential amino acids. This formulation is hereafter referred to as complete medium (CM). The same medium without serum, phenol red, and non-essential amino acids is referred to as incomplete medium (ICM).

Virus production medium (VPM; serum-free, containing 10 ng/mL EGF) was purchased from Gibco and supplemented with GlutaMax.

A 0.5 M EGTA stock solution (pH 8.0) was prepared in-house. Briefly, 19 g of EGTA (MW 380 g/mol) was dissolved in 90 mL of ddH2O. The pH was adjusted to 7.5-8.0 by gradually adding *>* 4 g of solid NaOH under mixing and gentle heating, with continuous pH monitoring. The final volume was adjusted to 100 mL.

FITC-dextran (average molecular weight 3000-5000 Da; Merck) was dissolved to 100 mg/mL in a 1:4 (v/v) mixture of DMSO and PBS (pH 7.4). A total of 15.234 mg of FITC-dextran was dissolved in 152 µL of solvent: first in 30.4 µL DMSO, followed by 121.6 µL PBS.

### 2.2 Transwell culture as barrier model

VERO E6 cells were obtained from the European Collection of Authenticated Cell Cultures (ECACC 85020206). Cultures were maintained in CM, and kept in humidified, 5% CO2 incubator at 37 °C.

For barrier transport experiments, the polycarbonate transwell inserts (0.6 cm^2^ area, 0.4 µm pore size, Merck Millipore) were equilibrated for at least 2 hours in 12 well plates (Sarstedt, Nümbrecht, Germany) with CM, then into each transwell insert 75 000 cells were seeded in 450 µL CM. At the same time, 1800 µL CM was added to the basolateral compartment. Cultures were grown for 5-7 days under normal culture conditions and were daily monitored by TEER measurements. Immediately before barrier transport measurements, transwell cell cultures and saturated cell-free transwell inserts were washed twice with PBS, then the apical and basolateral chambers were filled with 250 µL and 1200 µL ICM, respectively.

For cell free transport measurements transwell inserts were saturated with serum proteins by incubation in CM for 5-7 days. Unsaturated control transwell inserts were incubated in ICM.

### 2.3 Transepithelial electrical resistance (TEER)

Transepithelial electrical resistance (TEER) was measured using TEERScanner, a prototype automated scanning system (BioPhys-Concepts Kft, Budapest, Hungary, biophys-concepts.com) equipped with MERSSTX01 chopstick electrodes (Merck, Darmstadt, Germany) connected to a Millicell ERS-2 voltohmmeter (Merck, Darmstadt, Germany). During measurements, the 12-well plate was placed on a heated stage to maintain a constant temperature of 37 °C and transwell inserts were positioned within the wells using special transwell positioners to ensure consistent electrode placement.

Each measurement session involved two full scans of both the cell-free (blank) inserts and the inserts containing epithelial barriers, followed by a final third scan of the blank inserts. The entire scanning process for a 12-well plate was completed within approximately 5 minutes. To account for slow temporal drifts in baseline electrical resistance, interpolation was performed based on the resistance values obtained from the blank inserts. TEER values were calculated by subtracting the average resistance of the cell-free inserts from that of the cell-containing insert and normalizing the difference to the surface area of the transwell membrane, 60 mm^2^.

### 2.4 Live cell staining

For the visualization of cell monolayer integrity, various live cell stains were used. Cell-Tracker Green CMFDA dye (Thermo Fisher, C7025) was applied in 10 *µ*M concentration in the cell culture medium for 45-60 minutes to indicate intracellular esterase activity of live cells by conversion to green fluorescent compound in their cytoplasm. To visualize dead cells, monolayers were treated with red-fluorescent ethidium homodimer-1 (Thermo Fisher, L3224) at a concentration of 8 *µ*M for 20 minutes to indicate loss of plasma membrane integrity in dead cells where nuclei were labeled red. For visualization of all cell nuclei in the monolayer we used NucBlue (Hoechst 33342, Thermo Fisher, R37605) in 5 *µ*M concentration for 30 minutes resulting in blue fluorescent nuclear label.

### 2.5 Immunolabeling

Cells cultured on transwell membranes were fixed in 4% paraformaldehyde in PBS for 30 min, followed by permeabilization in 0.1% Triton X-100 in PBS for 10 min. Non-specific binding sites were blocked by incubation in CM for 1 h. Primary antibodies against beta-catenin (rabbit, polyclonal, Sigma C2206, 1/1000 dilution) or ZO-1 (rabbit, polyclonal, Sigma SAB3500301, 1/100 dilution) were applied for 2 h at room temperature then overnight at 4 ^*o*^C, followed by incubation in anti-rabbit secondary antibody conjugated with AlexaFluor-555 (Southern Biotech, 4030-32, 1/200 dilution) for 4 h at room temperature. All incubations were followed by triple washing steps in PBS for 30 min. Finally, the membranes were excised from the transwell insert devices and mounted on microscopic slides (Thermo Scientific) using mounting medium (Prolong Glass Antifade Mountant with NucBlue Stain, Invitrogen, P36981) to visualize cell nuclei.

### 2.6 Microscopy

Image acquisition was performed using a Zeiss Axio Observer Z1 inverted epifluorescent microscope with 5x EC Plan Neofluar, 10x Plan-Neofluar or 40x EC Plan-Neofluar objectives, and Zeiss Colibri illumination system with 365 nm, 470 nm and 555 nm LED modules and Zeiss HE25 filter set for fluorescence imaging. The microscope was equipped with a Zeiss AxioCam MRm CCD camera and a Marzhauser SCAN-IM powered stage. For multi-field mosaic image acquisition, stage positioning and focusing were controlled by Zeiss Axiovision 4.8 software. Images were processed using National Institute of Health (NIH) ImageJ software Schneider et al. [2012].

As described in Berta et al. [2021], time-lapse recordings were performed on a computer-controlled Leica DM IRB inverted microscope equipped with a Marzhauser SCAN-IM powered stage and a 10x HC PL Fluotar objective with 0.3 numerical aperture and 11 mm working distance. The microscope was coupled to Raspberry v2 camera. Cell cultures were kept at 37^*o*^C in humidified 5% CO_2_ atmosphere in a stage-top mini incubator during imaging. Phase contrast images of cells were collected consecutively every 10 minutes from each of the microscopic fields.

### 2.7 Segmentation of cell covered area

As described in Szeder et al. [2019] cell covered areas were identified by the presence of large local variability in brightness values. To detect cell occupied area a global threshold was applied to the local standard deviation of intensity on each image Wu et al. [1995], Neufeld et al. [2017]. The code used for segmentation and confluency calculation is available at http://github.com/aczirok/cellconfluency.

### 2.8 Automated transport sampling

As described in Tárnoki-Zách et al. [2023], Tarnoki-Zach et al. [2025] a prototype automated sampling system (BioPhys-Concepts, Budapest, Hungary, biophys-concepts.com) configured to utilize four separate tubing systems: two delivered the fresh medium while the other two removed the medium from the basolateral and apical chambers. The dosing unit used two syringe pumps and a fluid multiplexer to measure and inject 1.2 mL cooled medium into the tube connected to the incubator unit. A peristaltic pump is used to remove the medium from the device. Samples were collected in 2 mL Eppendorf tubes in the cooled fraction collector of the sampling system. Within the culture incubator, the fresh medium was kept in a heat exchanger for 30 minutes before being injected into the culture well. The culture was placed on a rotary shaker to ensure homogeneity of the basolateral compartment. The rotary shaker induced an intermittent rotary motion at a radius of 10 mm and a top rotational speed of 120 rpm. The time delay between the complete removal of the basolateral medium and the injection of fresh medium was less than 10 sec. Between measurements, the tubing network was cleaned and disinfected overnight using an automated procedure. First, all fluid handling volumes were filled for 20 minutes with 0.1% sodium hypochlorite. After the hypochlorite was removed, the tubing network was rinsed six times with sterile distilled water.

### 2.9 Spectroscopy

VIS absorption spectra were recorded with a Shimadzu UV-2101PC double beam spectrophotometer (Shimadzu, Kyoto, Japan) across the 370-750 nm spectral range at 2 nm spectral resolution, using distilled water as reference. Samples were equilibrated for at least 30 minutes at normal atmospheric CO_2_ conditions and loaded into 400 µL custom-fabricated cuvettes providing 1 cm optical path length. We used ICM in transport measurements to avoid absorption from phenol red and serum proteins. Both ICM and FITC exhibit absorption peaks below 550 nm (see Fig. 1). Thus, for baseline correction, a linear fit in the 550-750 nm spectral range was subtracted from the recorded spectra. Solvent-referenced spectra were obtained as the difference between the baseline corrected spectra of the sample and ICM.

**Figure 1:**
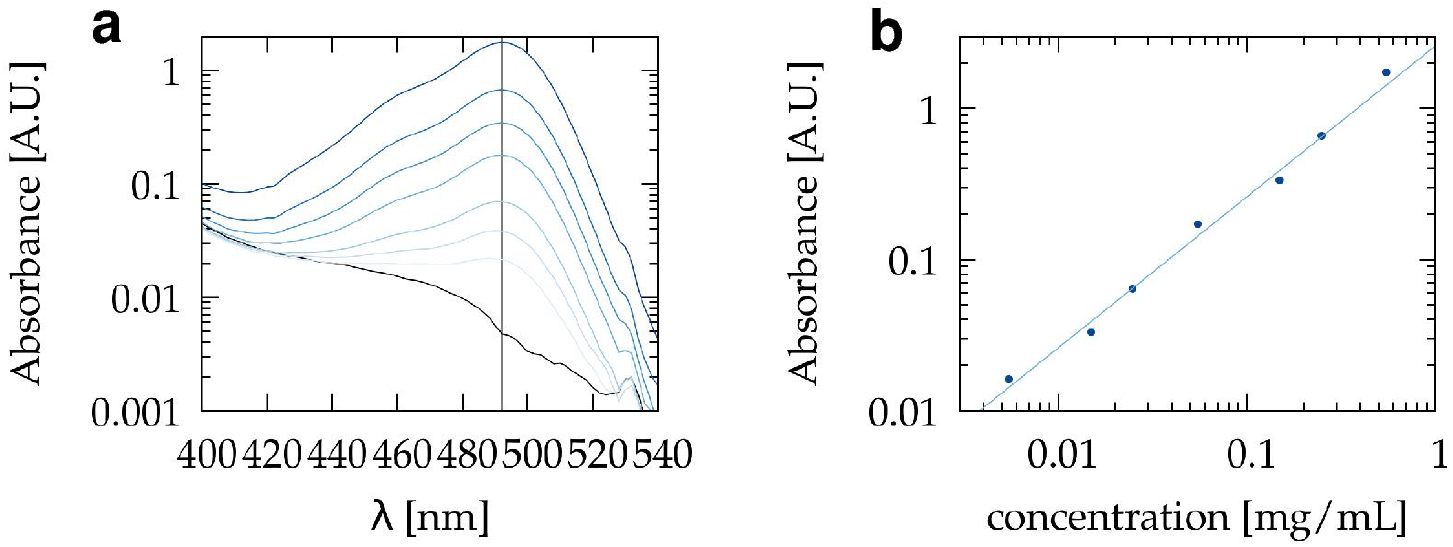
FITC-dextran calibration. (a) Baseline-corrected VIS absorption spectra of 0.005, 0.01, 0.02, 0.05, 0.1, 0.2, and 0.5 mg/ml solutions (higher concentrations shown in darker shades of blue) and ICM medium (black). (b) Peak absorbance values at 490 nm, plotted as concentration-dependent differences relative to ICM medium. A linear fit confirms the proportionality between concentration and absorbance within the experimental range.

### 2.10 Sampling correction

Measured analyte amounts in sequential samples often deviate from the true transport profile due to surface adsorption–desorption effects and residual droplet retention, which together can cause cross-sample contamination Tárnoki-Zách et al. [2023], Tarnoki-Zach et al. [2025]. To address these artifacts, we applied a linear recursive correction model, as detailed in Tarnoki-Zach et al. [2025]. The model parameters are obtained from cell-free calibration experiments, enabling the systematic removal of sampling-related distortions and yielding corrected concentration–time series suitable for downstream transport analysis.

### 2.11 Computational methods

Python codes were used for processing UV-VIS spectra, fitting model parameters and for the analysis of sample sequences. As described in detail in Tarnoki-Zach et al. [2025], data analysis has been performed using the scipy python module version 1.6.0. The analysis scripts are available at https://github.com/aczirok/barrier-transport-tools.

## 3 Results

### 3.1 Vero E6 cells form a barrier layer

To characterize barrier formation in Vero E6 monolayers, we analyzed parallel cultures seeded at different densities by phase-contrast videomicroscopy and by daily TEER measurements (Fig. 2). High-density cultures (160 × 10^3^ cells/cm^2^) were confluent from DIV (day *in vitro*) 1, whereas the lowest density (20 × 10^3^ cells/cm^2^) reached confluency by DIV 3. Across all seeding conditions, cells exhibited a characteristic cobblestone epithelial morphology and established stable contacts over time (Supplement Movie 1).

**Figure 2:**
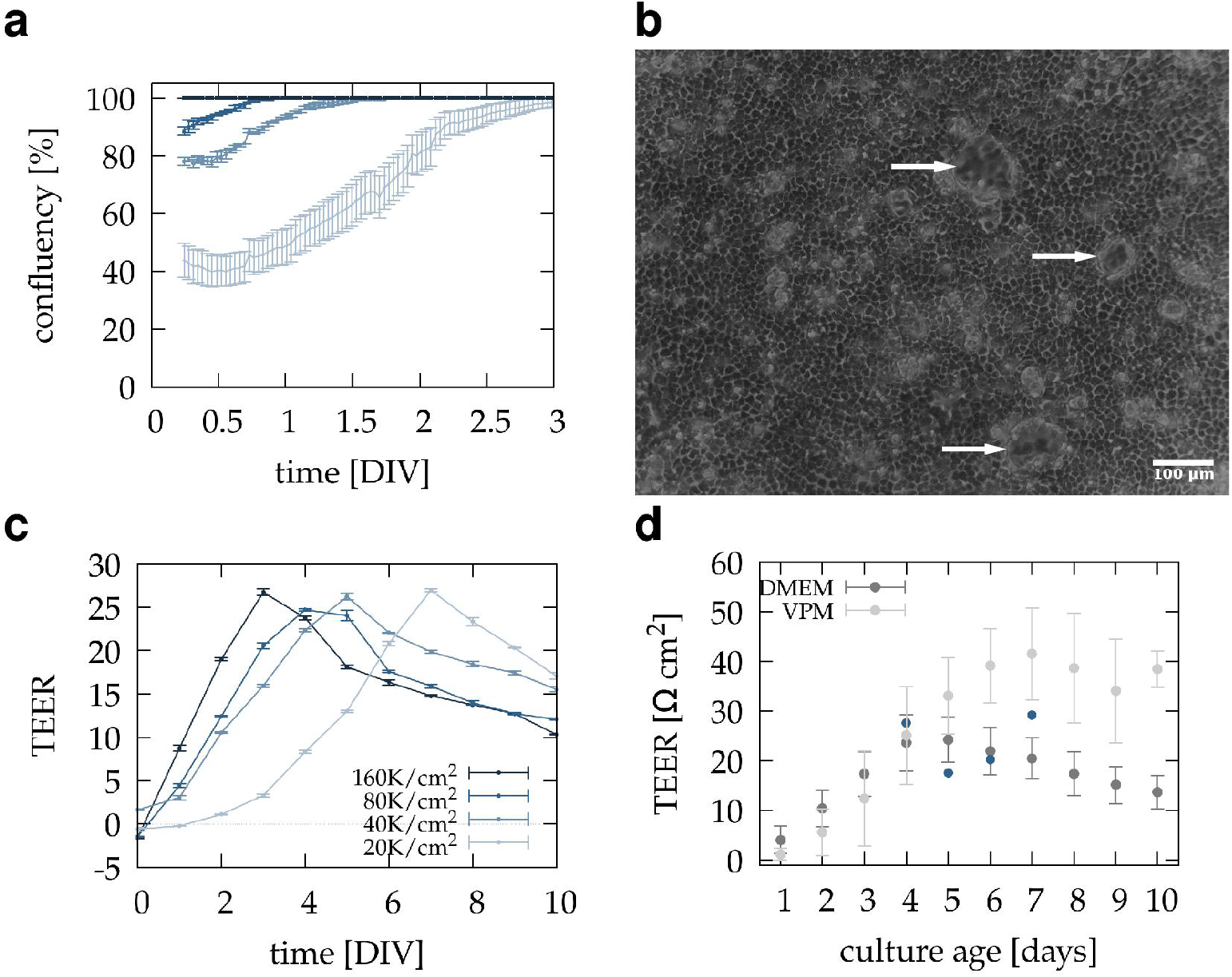
Barrier formation by Vero E6 monolayers. Vero E6 cells were seeded at densities of 20, 40, 80, and 160 × 10^3^ cells/cm^2^. Parallel cultures were followed either by time-lapse microscopy in standard culture dishes or by daily TEER measurements in transwell inserts. (a) Confluency quantified from videomicroscopy recordings (mean of 2–5 fields per condition). High-density (160 × 10^3^) cultures were confluent from DIV 1, while even the lowest density (20 × 10^3^) reached confluency by DIV 3. (b) Time-lapse images show dome formation (arrows) in confluent cultures, a morphological indicator of barrier maturation. (c) TEER values (mean of n = 2 technical replicates) demonstrated density-dependent kinetics, with all conditions reaching peak values between DIV 3 and DIV 7. (d) TEER development in cultures seeded at 80 × 10^3^ cells/cm^2^ and maintained either in serum-containing DMEM (n = 17) or virus production medium (VPM; n = 4). TEER rose at a similar rate in both media but reached a later and higher maximum in VPM. TEER values of cultures selected for FITC-dextran transport experiments are marked with blue symbols. Error bars represent standard deviation.

In confluent cultures, we observed prominent dome formation (Fig. 2b), a classical morphological indicator of epithelial maturation. Domes arise from directed ion and water transport across sealed junctions, which generates hydrostatic pressure beneath the monolayer and lifts the cell sheet from the substrate. Their appearance thus reflects both the acquisition of apical–basolateral polarity and the establishment of tight junctions. Dome formation is widely recognized as a hallmark of epithelial integrity *in vitro* and has been described in models such as MDCK cells Misfeldt et al. [1976], Cereijido et al. [1978].

TEER analysis of parallel cultures showed that functional maturation lagged behind morphological confluency (Fig. 2c). In CM, resistance values rose steadily to a peak of 30 Ω·cm^2^ between DIV 3 and DIV 7. Although the overall kinetics were similar across densities, cultures seeded at higher initial densities reached maximal resistance earlier.

To evaluate the effect of culture medium, we compared TEER development in cultures seeded at 80 × 10^3^ cells/cm^2^ and maintained either in CM or in VPM, a defined serum-free formulation used in virology (Fig. 2d). TEER rose at a comparable rate in both conditions but peaked later and at a higher value in VPM (40 Ω·cm^2^ at DIV 7). These results indicate that medium composition influences the extent of junctional maturation without altering the overall sequence of barrier development. Together, these findings demonstrate that Vero E6 monolayers reproducibly form confluent epithelial sheets with measurable barrier properties. The lag between confluency and maximal TEER likely reflects a maturation phase of intercellular junctions characteristic of epithelial barrier development.

### 3.2 Immunohistochemistry of Vero E6 monolayers

To assess monolayer integrity and junctional organization, Vero E6 cultures were examined by epifluorescence microscopy directly on intact transwell membranes. Nuclear (NucBlue) and cytoplasmic (CMFDA) dyes confirmed confluent coverage by day *in vitro* (DIV) 4, although cell density was moderately uneven, with accumulation near insert walls (Fig.3b). Viability staining with ethidium homodimer revealed a consistently low fraction of non-viable cells (*<* 1%), indicating robust survival during monolayer formation (Fig.3c).

To evaluate junctional maturation, cultures were fixed and immunostained for ZO-1, a scaffolding protein of tight junctions Itoh et al. [1997], and β-catenin, a key adherens junction component. By DIV 6, both markers exhibited clear cell–cell border localization (Fig. 3d–g), consistent with the establishment of mature intercellular contacts. Staining patterns appeared sharper and more continuous on glass coverslips than on polycarbonate membranes, reflecting differences in imaging conditions, but junctional labeling was robust under both substrates.

**Figure 3:**
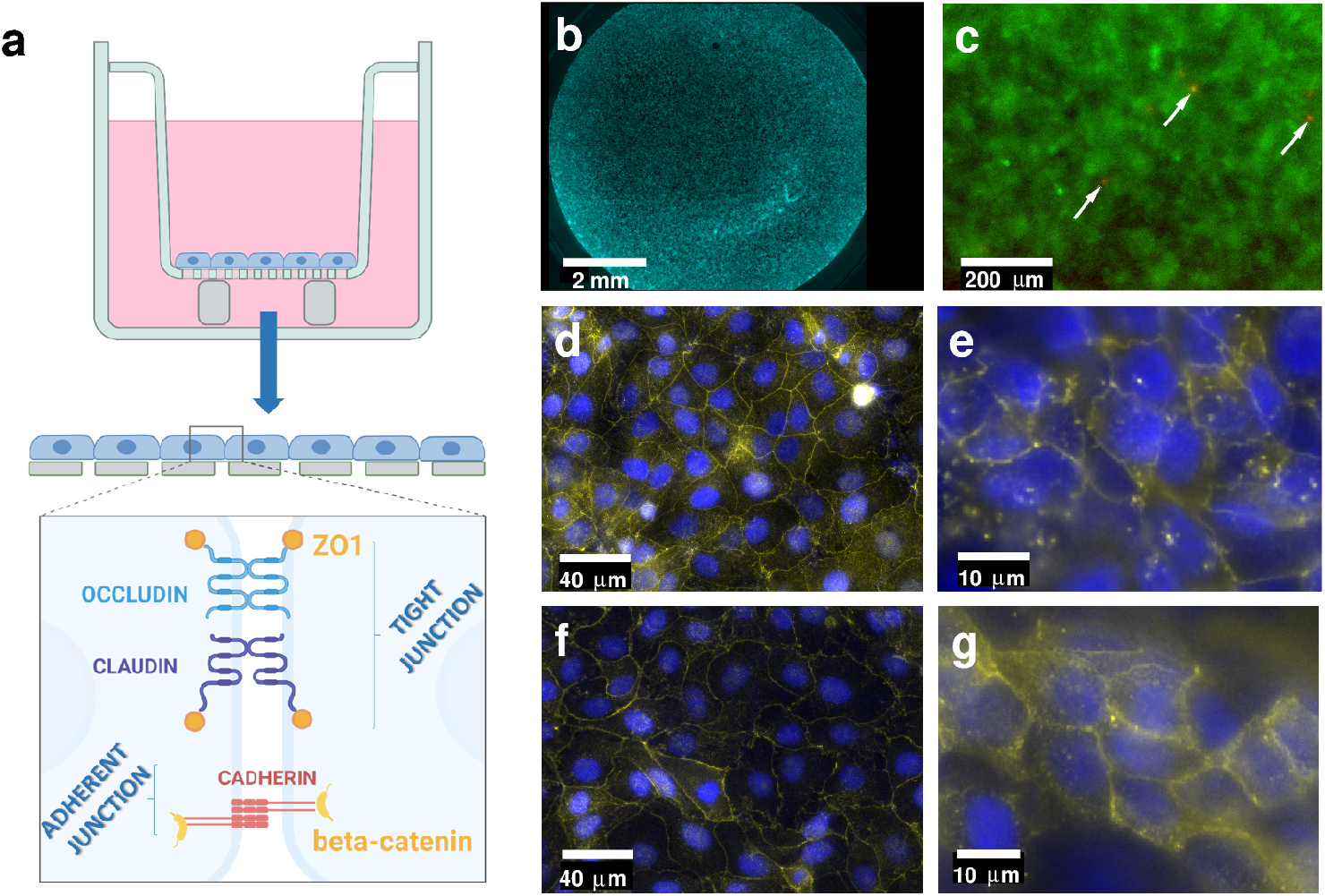
(a) Schematic representation of adherens and tight junction structures. (b) Nuclear staining with NucBlue shows uniform coverage of transwell membranes by day *in vitro* (DIV) 4 (5× objective). (c) Live/dead labeling with CMFDA (green, cytoplasm) and ethidium homodimer (red, nuclei of non-viable cells) demonstrates a continuous monolayer with a low fraction of dead cells (*<* 1%, arrows). (d-g) Immunofluorescence staining (yellow) shows junctional localization of the tight junction-associated proteins ZO-1 (d,e) and adherent junction protein β-catenin (f,g), in cultures grown on glass coverslips (d,f) and polycarbonate transwell membranes (e,g). Nuclei are counterstained with NucBlue (blue).

Together, these data demonstrate that Vero E6 monolayers form continuous, viable sheets with well-defined tight and adherens junctions by DIV 6. This structural evidence complements the electrical and transport measurements, supporting the conclusion that Vero E6 cells establish a functional epithelial barrier.

### 3.3 Reversible modulation of barrier integrity by EGTA

To probe the calcium dependence of Vero E6 barrier function, mature monolayers (DIV 5) were exposed to increasing concentrations of EGTA, a Ca^2+^chelator that disrupts tight junctions by sequestering extracellular calcium. Free calcium levels in DMEM under these conditions were estimated using a 1:1 stoichiometric binding model with a dissociation constant *K*_*d*_ = 2×10^−7^ M (Fig. 4a) Marks and Maxfield [1991], Mironov [2023]. Barrier integrity was dynamically assessed by recording TEER hourly during a 6-h EGTA treatment, with additional measurements obtained immediately before exposure and on the preceding day. After washout and replacement with calcium-containing medium, recovery was followed for several days at reduced temporal resolution.

**Figure 4:**
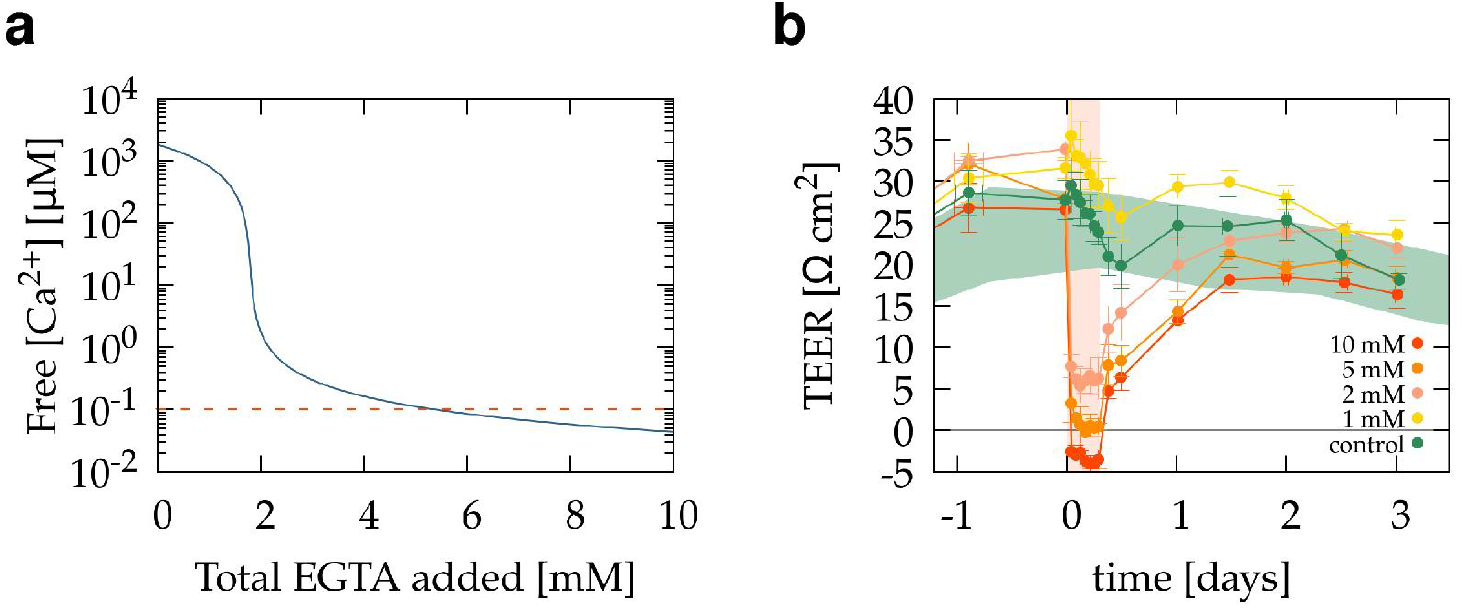
Reversible modulation of Vero E6 barrier integrity by EGTA. (a) Estimated free Ca^2+^concentrations in DMEM as a function of added EGTA, calculated assuming 1:1 binding with *K*_*d*_ = 2 × 10^−7^ M. (b) TEER of mature Vero E6 monolayers (DIV 5) during exposure to 1–10 mM EGTA. A slight TEER increase occurred with 1 mM EGTA, whereas 2–10 mM induced rapid, dose-dependent, and reversible decreases. The peach-shaded area indicates the 6 h EGTA treatment period. Time is shown in days relative to EGTA addition (*t* = 0). Following washout, TEER progressively recovered. Data represent mean ± SD from at least two inserts per condition, pooled from two independent experiments. Horizontal error bars denote variability in measurement timing. The green band indicates the TEER range (mean ± SD) of untreated cultures over time (n=17).

As shown in Figure 4b, 1 mM EGTA caused a modest rise in TEER, most likely reflecting reduced medium conductivity rather than genuine tightening of junctions. In contrast, 2–10 mM EGTA induced rapid, dose-dependent TEER decreases, with values approaching or dropping below those of blank inserts. The largest reduction was observed with 10 mM EGTA, consistent with combined effects of junctional disruption and increased ionic strength of the medium.

Upon reintroduction of calcium, TEER values progressively recovered, confirming that the effect was largely reversible. Cultures treated with 2 mM EGTA regained baseline resistance within 24 h, whereas recovery from 5–10 mM required up to 36 h. These findings demonstrate that Vero E6 tight junctions are functionally dependent on extracellular calcium and capable of restoring barrier integrity following chemical perturbation.

### 3.4 Dextran permeability

To evaluate the barrier properties of Vero E6 monolayers, we measured apical-to-basolateral transport of 4 kDa FITC-dextran, a paracellular tracer that does not cross intact tight junctions or enter cells. Vero E6 cultures were maintained on transwell membranes in CM until TEER reached a plateau (typically DIV 5), after which inserts were placed into a Millitransflow fluid-handling system (BioPhys-Concepts, Budapest). The system collected basolateral samples automatically at 30-min intervals for 6 h, with two baseline samples taken before apical replacement with FITC-dextran solution (0.5 mg/mL). FITC-dextran concentrations were determined by UV–VIS spectrophotometry.

As shown in Figure 5, cell-free control membranes allowed rapid tracer accumulation in the basolateral compartment, peaking in the first sample. In contrast, Vero E6 monolayers markedly restricted FITC-dextran passage, with basolateral levels remaining near background throughout the experiment. By the end of the assay, nearly all tracer remained in the apical compartment, confirming that the monolayer effectively blocked paracellular diffusion.

**Figure 5:**
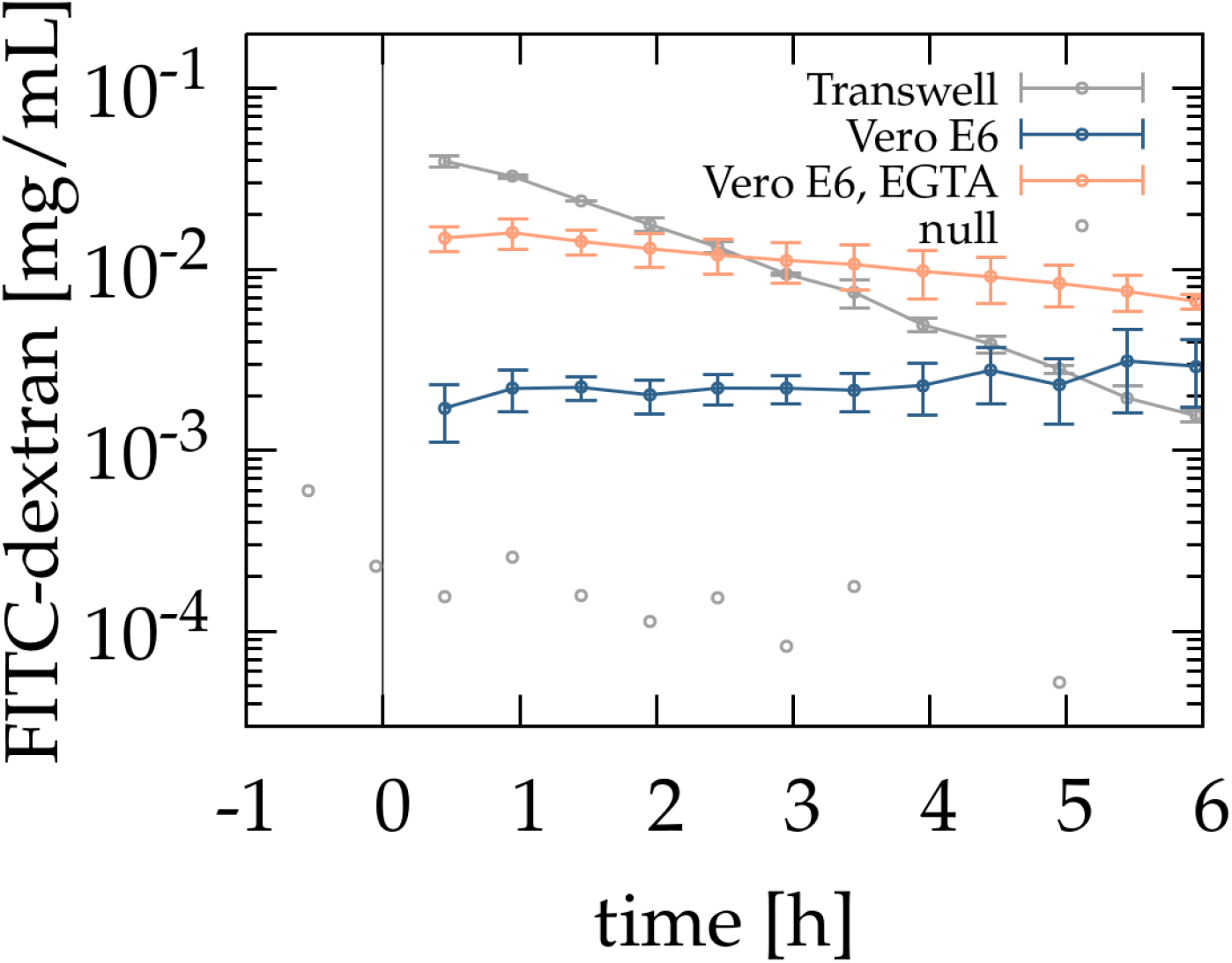
Vero E6 cells form a barrier that restricts FITC-dextran paracellular permeability. Apical-to-basolateral transport of 4 kDa FITC-dextran was monitored over 7 h with automated 30-min sampling. Cell-free membranes (grey dots) showed rapid tracer accumulation, whereas membranes sealed with silicone grease (grey open circles) served as impermeable controls. Vero E6 monolayers (blue symbols) reduced FITC-dextran permeability by more than an order of magnitude compared to cell-free inserts, maintaining basolateral concentrations near background. Barrier disruption with 2 mM EGTA (orange symbols) significantly increased transport, confirming calcium-dependent regulation of paracellular permeability. Data represent means ± SD from three independent experiments for each condition.

Apparent permeability coefficients (P_app_), calculated from cumulative transport data, were typically 3 − 5 × 10^−6^ cm/s. These values are comparable to those reported for established epithelial and endothelial barrier models Ma et al. [2007], Francia et al. [2018], indicating that Vero E6 cultures form physiologically relevant barriers suitable for transport studies.

To assess calcium dependence, transport assays were repeated wiin the presence of 2 mM EGTA, a non-cytotoxic concentration that partially disrupts tight junctions. Under these conditions, permeability increased nearly an order of magnitude (Fig.5), confirming that barrier restriction is calcium-dependent. TEER measurements obtained at the end of transport assays matched the corresponding TEER calibration data (Fig.4), validating that the assay itself did not compromise barrier function. Together, these findings demonstrate that Vero E6 monolayers form continuous, low-permeability barriers whose function can be reversibly modulated by extracellular calcium, supporting their utility as a model for mechanistic studies and compound screening.

## 4 Discussion

### 4.1 Vero E6 as a hybrid model for infection and transport studies

By integrating barrier function measurements (e.g., TEER, macromolecular permeability, and junctional protein localization) with traditional virology assays, researchers can move beyond viral replication alone and assess how infection affects host tissue architecture and defense Linfield et al. [2021], Mamana et al. [2023], Ruan et al. [2020]. This approach also enhances our ability to model drug penetration Shirsath et al. [2024], Rahman et al. [2025], immune cell trafficking Yonker et al. [2017], Os et al. [2023], and co-infection dynamics Walch and Broz [2025], especially in systems mimicking mucosal surfaces, blood-brain barriers, or lung alveoli. Furthermore, barrier models are useful for studying viral dissemination mechanisms, such as transcytosis, paracellular leakage, or junctional remodeling, which cannot be adequately captured in non-polarized or suspension cell systems Knyazev et al. [2021], Tugizov et al. [2013], Dong et al. [2020], Ding et al. [2022], Day et al. [2022], Chapuy-Regaud et al. [2022]. In the context of high-impact pathogens like SARS-CoV-2, which target epithelial surfaces and are capable of disrupting barrier integrity Short et al. [2016], Tugizov [2021], Ding et al. [2022], Linfield et al. [2021], combining barrier biology with virology becomes especially relevant.

In this study, we systematically evaluated the barrier-forming properties of Vero E6 cells, a widely used cell line in virology, but comparatively understudied in the context of epithelial barrier biology. Our results show that Vero E6 cells seeded at appropriate densities reliably form confluent monolayers that develop measurable transepithelial electrical resistance (TEER), limit the paracellular passage of macromolecules such as FITC-dextran, and express key junctional proteins including ZO-1 and β-catenin. Moreover, these monolayers exhibit reversible barrier disruption in response to calcium chelation by EGTA, confirming the dynamic regulation of their junctional complexes.

### 4.2 Barrier formation and dynamics in Vero E6 cultures

Our time-resolved TEER recordings and videomicroscopy data reveal that monolayer formation and barrier maturation follow a density- and time-dependent trajectory. Cultures seeded at moderate densities reach confluence by DIV 2–4, while TEER peaks later, typically around DIV 5, with values in the range of 30-45 Ω·cm^2^ depending on the culture medium. Although these TEER values are modest compared to classical tight epithelial barriers such as Caco-2 or MDCK II, they fall within the range commonly observed in endothelial monolayers. Importantly, FITC-dextran transport measurements revealed a paracellular permeability (Papp) of approximately 3.7× 10^−6^ cm/s, a value comparable to established endothelial barriers including human umbilical vein endothelial cells (HU-VECs) and corneal endothelium Ma et al. [2007], Francia et al. [2018]. This underscores the importance of considering permeability assays in addition to TEER when evaluating barrier function: even cultures with relatively low resistance can exhibit physiologically relevant selectivity for macromolecular transport.

The formation of continuous monolayers and the detection of key tight (ZO-1) and adherens (β-catenin) junction components further support the notion that Vero E6 cultures develop a structurally and functionally competent epithelial interface. While they may not model tight barriers in the classical sense, their moderate resistance and selective permeability make them highly relevant for studies where barrier modulation and pathogen interaction are of interest.

### 4.3 Sensitivity to EGTA reveals calcium-dependence and reversible modulation

EGTA treatment induced rapid and reversible decreases in TEER, consistent with the calcium dependence of tight junction integrity Rothen-Rutishauser et al. [2002]. Dose-dependent TEER reductions were observed starting from 2 mM EGTA, with recovery kinetics strongly affected by the concentration. These effects mirror findings in other epithelial systems, reinforcing the idea that Vero E6 tight junctions are functionally regulated by extracellular Ca^2+^Tria et al. [2013], Panou et al. [2023]. The associated increase in FITC-dextran permeability confirms that paracellular transport is sensitive to junctional modulation, and that EGTA treatment can serve as a controlled experimental perturbation to probe barrier plasticity in Vero E6 cultures.

### 4.4 Limitations and future directions

Although Vero E6 cells form functional monolayers, their baseline TEER values are modest, and their tight junction architecture may differ from highly polarized intestinal or renal epithelia. Moreover, as a non-human, kidney-derived cell line, Vero E6 cells may not fully replicate tissue-specific responses of human epithelia. Nonetheless, their virus-permissive phenotype, ease of handling, and barrier-like behavior make them valuable for compound screening, infection models, and studies linking viral effects to barrier function. Future work could investigate the transcriptional regulation of junctional proteins, compare the barrier properties of Vero E6 with other permissive cell lines, and assess how virus infection alters barrier integrity in this system. Coupling live-virus infection with real-time TEER and permeability assays may yield new insights into how pathogens compromise epithelial defenses.

